# Hedonic experiences emerge from an orchestrated balance of synergistic and redundant information processing

**DOI:** 10.1101/2025.10.05.680521

**Authors:** M. Kathofer, P.A.M. Mediano, M. Spies, A. Liardi, G. Dörl, P. Stöhrmann, C. Schmidt, E. Briem, G. Schlosser, B. Eggerstorfer, M. Klöbl, H. Leder, R. Lanzenberger, J.S. Crone

## Abstract

Ketamine exerts rapid-acting, pro-hedonic effects, yet its precise mechanism remains elusive. Here, we present behavioral and fMRI data from a randomized, placebo-controlled crossover study in 38 healthy participants investigating ketamine’s sub-acute effects on multivariate information-processing during music-evoked peak hedonic experiences. Leveraging information-theoretical measures, our findings indicate that hedonic experiences depend on a distinct global (as measured by O-Information) and local (as measured by integrated information) balance between redundant – information shared across nodes – and synergistic – information emerging from joint interactions – processes. As hedonic intensity rises, neural dynamics shift toward greater synergy; with the one exception of a deliberate increase in redundancy particularly for key sensory information to ensure reliable transmission of and access to critical external information for subsequent hedonic processing. In contrast, ketamine’s sub-acute pro-hedonic effects arise potentially from enhancing redundant dynamics at rest, boosting the brain’s capability to robustly represent and access critical internal information, and thus, fostering an environment optimized to amplify the phenomenological hedonic experience, while simultaneously allowing for more efficient information integration.

## Introduction

The hedonic experience is a fundamental aspect of mental health, and its disruption, called anhedonia, is a core symptom of many psychiatric disorders, including depression, schizophrenia, and substance use disorder^1–3^. Addressing anhedonic symptoms specifically is critical for improving treatment outcomes^4–7^ and offers the potential for more effective therapies, as emphasized by the National Institute of Mental Health (NIMH), which declares anhedonia as a key focus of its research domain criteria framework^8^. One particularly promising intervention for anhedonia is ketamine, which garnered significant attention for its rapid-acting pro-hedonic effects over the past decade^9^. Administered at a subanesthetic dose, ketamine alleviates depressive^10–12^ and anhedonic^13,14^ symptoms within hours. However, the neural mechanisms underlying these effects remain elusive, as much of the existing research relies on post-hoc self-report questionnaires that fail to capture real-time changes in hedonic processing. Since the hedonic experience is thought to emerge from the orchestrated interplay of various cortical and subcortical regions^15–17,see^ ^18^ ^for^ ^a^ ^review^, information theory, a framework that conceptualizes the brain as a multivariate information processing system, serves as a particularly powerful tool for studying these interactions. Thus, combining real-time assessments of hedonic processing with insights from information theory offers a promising avenue to deepen our understanding of the hedonic experience itself and illuminate how ketamine shapes neural dynamics to enhance hedonic experiences.

Higher-order information integration, arising from interactions that span more than two brain regions, is a key feature of brain function. Two complementary forms of integration – synergy and redundancy – offer valuable insights into these multivariate dynamics^19^. Synergy refers to information that emerges solely from the joint activity of multiple regions and cannot be found in any single region alone, thus facilitating more efficient information processing^20^. However, since it depends on the coordinated contribution of these regions, synergistic information is fragile and can be easily lost if even one contributing region is disrupted. In contrast, redundancy indicates information that is duplicated across regions, ensuring robust representation of critical information. Both synergy and redundancy play crucial roles in behavioral and cognitive task performance^21,22^, with evidence suggesting that an optimal balance between the two is necessary for effective functioning. Together, they reflect a fundamental trade-off in the brain’s integration strategies, balancing the adaptability of emergent information processing with the reliability of robust information representation.

Recent methodological advances have begun to make these higher-order interactions accessible to empirical investigation, each with their own distinct strengths and limitations^23–25^. One such approach, known as *information about organizational structure* (O-Information)^23^, provides a simple way to assess whether the information within a large set of brain regions is dominated by redundancy (positive O-Information) or synergy (negative O-Information). While O-information does not explicitly separate these components, it summarizes their balance, providing insights into the dominant mode of multivariate interdependence in a system. On the other hand, *whole-minus-sum integrated information* (Φ^WMS^)^26^ can be utilized to investigate how integrative hedonic processes evolve over time. More specifically, Φ^WMS^ examines how brain regions coevolve over time by assessing whether the joint activity of the regions (the “whole”) is better at predicting their combined future than the individual regions (the “parts”) are at predicting their own. While Φ^WMS^ provides crucial insights into integration dynamics, the recently developed integrated information decomposition (ΦID) framework^27^ offers a more intuitive interpretation by allowing Φ^WMS^ to be decomposed into its two key components: Φ^R^ and redundancy (rtr) (Φ^WMS^ = Φ^R^ – rtr). Thereby, Φ^R^ represents a revised measure of integrated information^28^, capturing all synergistic temporal processes as well as information that is transferred from one region to another. In contrast, rtr quantifies whether shared information between regions persists over time, reflecting redundant information storage across brain areas. Thus, positive Φ^WMS^ values indicate that synergistic and transfer processes dominate the temporal dynamics, while negative values suggest that redundant processes outweigh synergistic ones. By disentangling these components, ΦID offers a deeper understanding of the mechanisms driving temporal information integration during neural processing. Leveraging these complementary measures (O-Information, Φ^WMS^, and its constituents, Φ^R^ and rtr) provides a unique opportunity to overcome the limitations of each measure individually, shedding light on how distributed brain areas coordinate to give rise to the hedonic experience and how these processes may be modulated by interventions such as ketamine.

To do so, the present study recorded fMRI data during a hedonic task involving self-selected music, a potent and personalized stimulus capable of reliably inducing peak hedonic experiences^29–32^. While ketamine’s effects are typically studied in clinical populations suffering from anhedonia, this study focused on a healthy population to minimize confounds such as comorbidities and prior treatment effects. This allowed us to probe the following 4 hypotheses regarding information dynamics underlying the emergence of a hedonic experience in general:

H1: We hypothesized that ketamine would exert measurable pro-hedonic effects in the healthy population.

H2: Given the inherently emergent nature of hedonic experiences – which rely on the integration of both external (e.g., sensory perception) and internal (e.g., cognitive and affective) information^32^ – we hypothesized that hedonia arises from enhanced synergistic information integration. Specifically, we expected to observe shifts in both O-Information and Φ^WMS^ measures toward greater synergy during hedonic states.

H3: Furthermore, we hypothesized that ketamine would generally shift neural information dynamics toward greater synergistic integration, promoting a neural state more conducive to the emergence of intense hedonic experiences.

H4: Finally, because hedonic experiences depend on the sustained integration of internal and external inputs over time, we hypothesized that hedonic states would be characterized by a tendency to maintain a highly synergetic state over time.

## Results

### Ketamine increases hedonic response across multiple dimensions

To reliably induce a range of hedonic experiences in a more ecologically valid setting, we designed an individualized hedonic task (Figure 1A) using self-selected music – a dynamic, real-time stimulation known to elicit strong affective responses^29–32^. During this task, each participant (*n* = 38) rated their hedonic responses to 70-second excerpts of 10 songs they found highly pleasurable and 10 songs they deemed neutral (initial judgment) on three dimensions: valence (emotional value), moving (being emotionally moved), and chills (perceived chills). The task was performed during fMRI in two sessions: four hours after ketamine (0.5 mg/kg) and four hours after placebo (0.9% saline) administration (Figure 1B). Each session included eight interleaved 30-second baseline segments to evaluate ketamine’s sub-acute effects on intrinsic neural dynamics at rest throughout the task.

**Figure 1.**
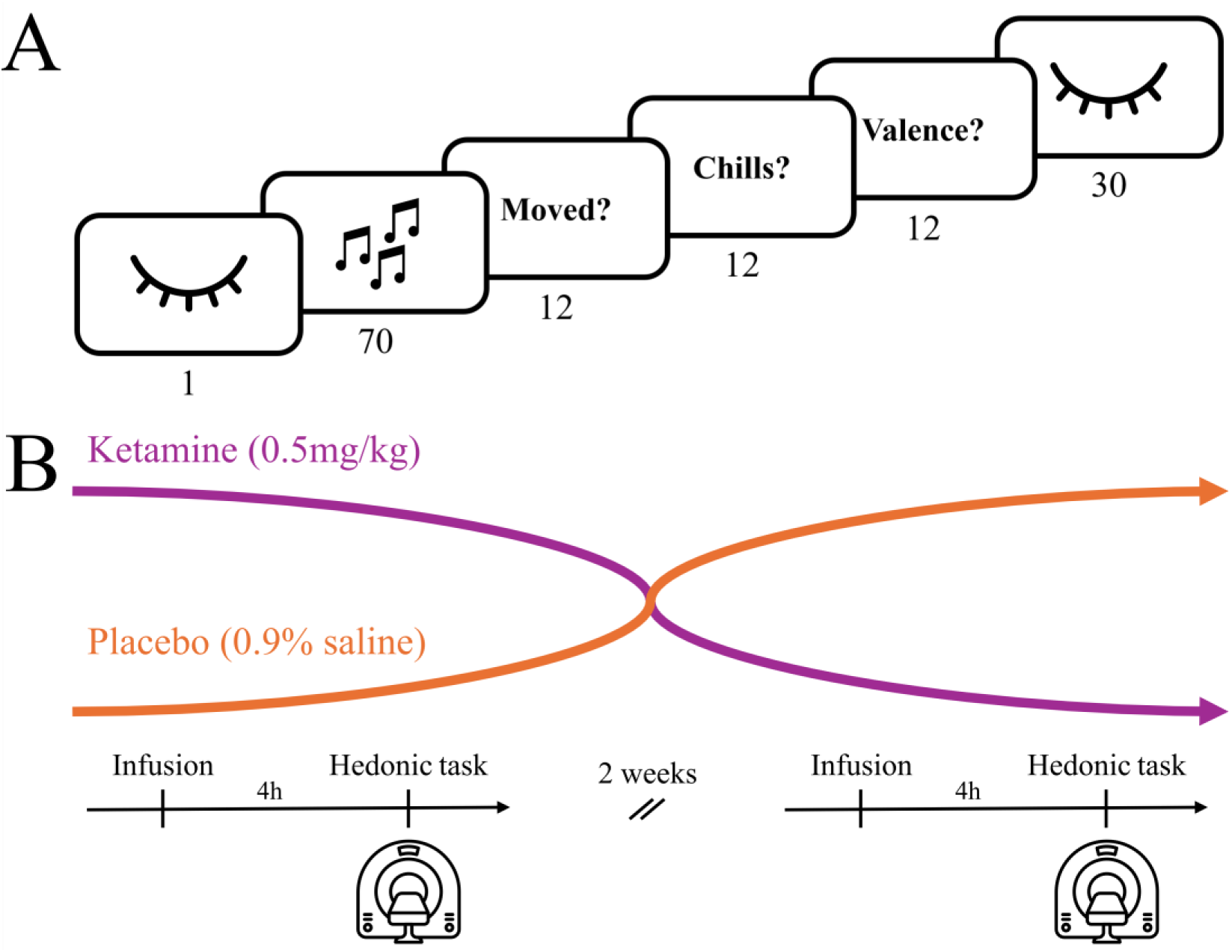
Hedonic task and study design. (A) Example of one music trial during the hedonic task. With eyes closed, participants listened to a 70-second excerpt from either one of their self-selected 10 most pleasurable songs or 10 neutral songs. After each excerpt, they rated the quality of the evoked hedonic experience along three dimensions: moving, chills, and valence. The task was divided into two runs, each consisting of 9-12 music trials and four interleaved 30-second eyes-closed baseline (resting) segments. Numbers beneath pictograms indicate the duration of each section in seconds. (B) 38 healthy participants (18–55 years) completed a randomized, single-blind, placebo-controlled, cross-over study. Each underwent two fMRI sessions, separated by at least two weeks. Four hours before scanning, participants received either a placebo or 0.5 mg/kg racemic ketamine (40-min infusion), with arm allocation randomized. During each session, they performed the hedonic task.

Behavioral results confirmed successful hedonic manipulation across all tested dimensions: positive songs were considerably more likely than neutral songs to evoke strong responses in valence (Odds Ratio [OR] = 116.08, p < 0.01, Bonferroni-adjusted α = 0.016), moving (OR = 219.68, p < 0.01), and chills (OR = 114.05, p < 0.01) (see Tables S1-S3). As hypothesized (H1), ketamine further enhanced hedonic responses across all three tested dimensions, increasing valence (OR = 1.36, p < 0.01), amplifying moving (OR = 1.31, p < 0.01), and markedly enhancing chills (OR = 1.60, p < 0.01). Based on prior model comparisons, no interaction term between treatment (ketamine vs. placebo) and initial judgment (positive vs. neutral) was included in the mixed ordinal regression, suggesting that ketamine promotes a richer and more complex affective and physiological response regardless of the stimulus’ initial emotional connotation.

### Music perception and intense hedonic experiences shift global information processing toward synergy

Having established that ketamine does indeed increase all dimensions of the hedonic experience in healthy individuals, we turned to the question of whether hedonic experiences shift the global balance in information integration toward synergy. To do so, preprocessed BOLD data (*n* = 32) was parcellated into 116 cortical^33^ and subcortical^34^ regions, and averaged O-Information values for each music trial and baseline segment were calculated, yielding one value per music trial and baseline segment per subject.

Importantly, our hypothesis (H2) is based on an implicit assumption: when engaging with a dynamical, sensory-rich stimulus such as music, the brain should already naturally shift toward more synergistic dynamics compared to rest, as it requires continuous integration across time. To test this, we fitted a linear mixed-effects model with the O-Information values of the music trial and baseline segments as dependent variables, including fixed-effects regressors for music perception (music vs. baseline), treatment (ketamine vs. placebo), and a random intercept for each participant. Three key findings emerged. First, the brain in its natural state exhibited a highly redundant organization during rest (O-Information = 32.26 nats; see Table S4). Second, during music processing, O-Information shifted significantly toward synergy compared to baseline (standardized beta (*β_stand_)* = −0.11, *SE* = 0.04, *t*(1748) = −2.46, *p* = 0.01, Bonferroni-adjusted α = 0.025). Third, ketamine did not affect global information dynamics (*β_stand_* = 0.02, *SE* = 0.04, *t*(1748) = 0.49, *p* = 0.62), which was further validated using exclusively the baseline segments (see Table S5).

To examine whether the intensity of subjective hedonic experiences during music stimulation was associated with changes in global neural dynamics, we fitted another linear mixed-effects model, including treatment (ketamine vs placebo) and hedonic intensity (moving rating) as predictors, with a random intercept for participants. Consistent with our hypothesis (H2), we found that more intense hedonic experiences were associated with a greater shift toward global synergistic processing (*β_stand_* = - 0.07, *SE* = 0.02, *t*(1230.9) = −3.02, *p* < 0.01) (see Table S6). Again, ketamine did not significantly alter global information dynamics during music perception (*β_stand_* = 0.05, *SE* = 0.05, *t*(1222.2) = 1.10, *p* = 0.27), suggesting that despite global dynamics shifting in response to the phenomenological experience, the global dynamics during music processing remained unaffected by the neuromodulatory effects of ketamine.

### Ketamine, at rest, induces widespread shifts toward redundancy, whereas the hedonic experience induces local shifts toward both synergy and redundancy

While O-Information provides valuable insights into higher-order information dynamics of large-scale systems, it does not explicitly capture how past neural activity shapes future dynamics – a critical aspect of music-evoked hedonic processing^32^. To address this, we used Φ^WMS^, a measure of dynamic information integration, to track changes in neural integration over time and identify central hubs involved in hedonic processing. Time-delayed mutual information terms were calculated for each run and then averaged within each trial window to produce mean Φ^WMS^ matrices^35^, reflecting time-averaged information dynamics across all pairs of brain regions. To identify changes in neural processing associated with the subjective intensity of the hedonic experience, we analyzed shifts in Φ^WMS^ using a modified version of the network-based statistic (NBS) algorithm^36,37^, capable of detecting clusters with both positive (synergy-increasing) and negative (redundancy-increasing) edges simultaneously.

A highly localized cluster (*p*_fwer_ < 0.01) associated with the main effect of the subjective hedonic experience was identified, comprising 109 positive (Φ^WMS+^) and 68 negative (Φ^WMS-^) edges (see hedonic cluster in Figure 2A). More specifically, interactions shifting toward greater synergy increased markedly within and between the parietal lobe, frontal, and temporal regions, as the intensity of the hedonic experience grew. Strikingly, over one-third (38 of 109) of the synergy-increasing edges involved the insula, pointing to its central role as a hub of interoceptive integrative processing^38,39^. In contrast, a majority of the redundancy-increasing edges emerged between subcortical and frontal structures. The most prominent hub in this subnetwork was the globus pallidus, appearing in 19% of all redundancy-increasing edges. This subcortical region increased redundant, that is, stable interactions with a broad range of areas, including the nucleus accumbens, putamen, insula, and orbitofrontal cortex. Notably, as the hedonic experience intensified, joint information dynamics between the auditory cortex and regions such as the insula, prefrontal cortex, and thalamus, became increasingly redundant, suggesting a more stable relay of auditory content during stronger subjective experiences.

**Figure 2:**
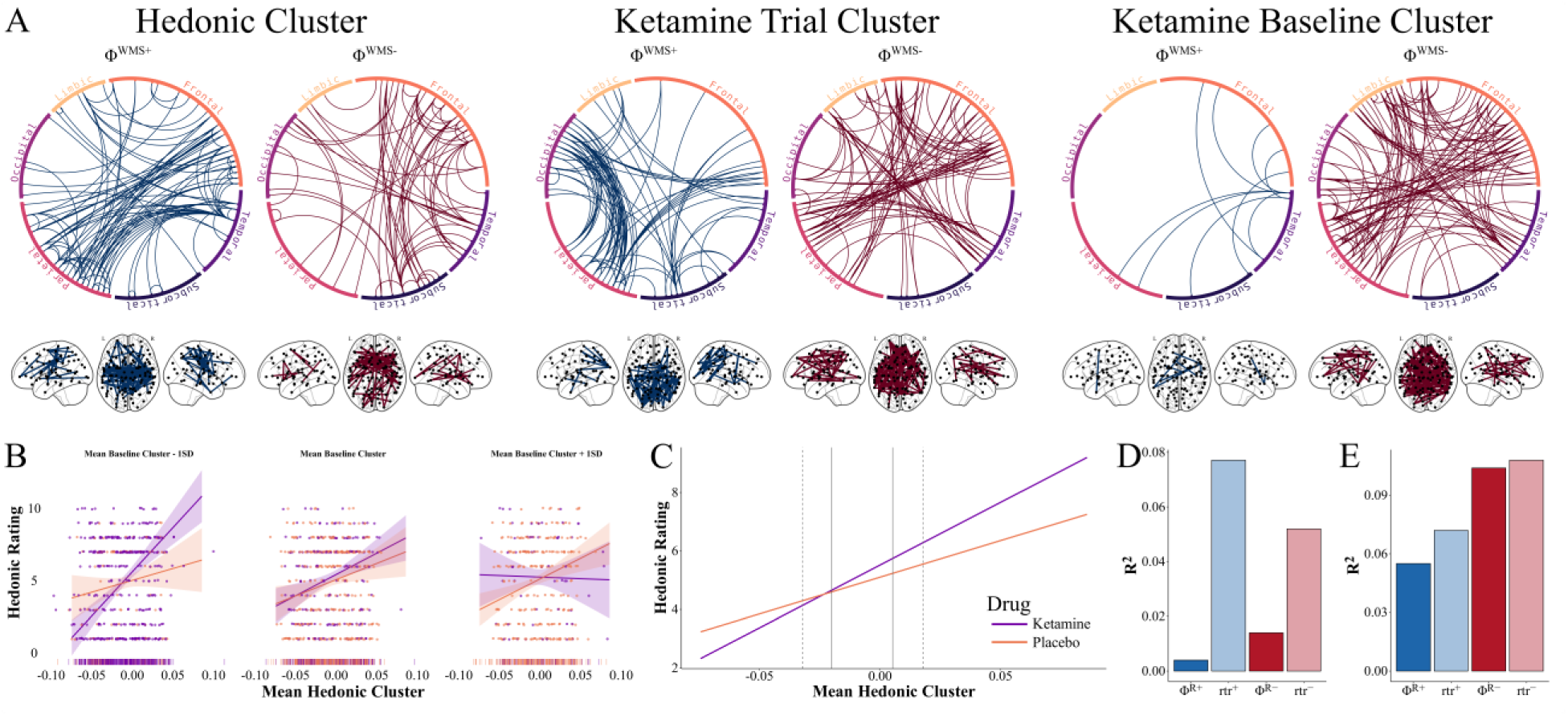
Hedonic experiences induce local shifts toward both synergy and redundancy. (A) Significant NBS clusters are displayed with increasing edges (Φ^WMS+^) and decreasing edges (Φ^WMS-^) for all pairs of regions separately. Edges are visualized both using a chord diagram (upper panel) and in a glass brain (lower panel) for each of the cluster solutions: the hedonic cluster, the ketamine trial cluster, and the ketamine baseline cluster. (B) The three panels illustrate the identified three-way interaction between treatment (ketamine vs placebo), the average activity of the hedonic cluster, and the average activity of the baseline ketamine cluster (both measured in nats) in predicting moving experiences. The interaction is visualized across ±1 standard deviation (SD) of the baseline ketamine cluster average. Below, carpet plots display the distribution of observations across baseline ketamine cluster values for each treatment. Due to sparse data at the extreme ends (i.e., low baseline values under placebo and high baseline values under ketamine), the confidence intervals of the inference lines are wider, complicating visual inference. (C) To address this, this panel presents a clearer visualization of the interaction: predicted values of moving experience are plotted across the full range of the hedonic cluster average, separately for two extracted levels of the baseline ketamine cluster (−1 SD under ketamine, +1 SD under placebo). Vertical solid grey lines indicate the ±1 SD range of the hedonic cluster values; dashed grey lines indicate the ±2 SD boundaries. (D) Dominance analysis for the hedonic cluster. Bars represent the average explained variance of each predictor. (E) Dominance analysis for the drug logistic model.

No significant interaction was observed between ketamine administration and moving ratings on neural integration. However, NBS recovered one significant cluster reflecting changes in information dynamics specifically attributed to the main effect of ketamine during music perception (79 positive edges; 92 negative edges; *p*_fwer_ < 0.01). This cluster was characterized by a marked shift toward greater redundancy (Φ^WMS-^) between frontal and parietal regions, alongside a shift toward greater synergy (Φ^WMS+^) between parietal and occipital areas (see ketamine trial cluster in Figure 2A). The most prominent hub for Φ^WMS+^ was the primary sensory cortex, involved in 34% of the synergy-increasing edges. In contrast, the most frequent hub associated with a shift toward greater redundancy (Φ^WMS-^) was the right paracingulate cortex (14%).

Given that ketamine’s pro-hedonic effects often persist for up to a week after administration and appear largely independent of the initial valence of the stimulus, enhancing a broad range of experiences, as supported by both prior literature^40–42^ and our behavioral findings, we hypothesized that ketamine induces sustained changes in intrinsic neural dynamics that support its therapeutic action. Thus, testing ketamine’s effects during music stimulation may be suboptimal for uncovering these mechanisms.

Instead, we conjectured that alterations in the absence of stimulation, that is, during rest, should show a shift toward greater synergistic processing, and that these fundamental functional alterations may prime the brain to process subsequent stimuli in a way that enhances the hedonic experience (H3). To test this, we examined ketamine’s effects on resting dynamics by applying the same NBS analysis to the mean Φ^WMS^ matrices from the interleaved baseline segments, thereby isolating information integration patterns independent of stimulation (music input). A single significant cluster with a prominent pattern emerged (*p*_fwer_ < 0.01); 93% of the edges following ketamine administration shifted toward greater redundancy (Φ^WMS-^) with 122 negative and only 9 positive edges. This shift toward greater redundancy was most pronounced between frontal, parietal, and subcortical regions (see ketamine baseline cluster in Figure 2A).

### Ketamine fundamentally reshapes associations between neural integration and subjective experience

Given these findings, we explored the idea that ketamine’s pro-hedonic effects may stem from its ability to shift resting dynamics toward greater redundancy, enhancing the brain’s ability to reliably access critical internal representations. Generally, hedonic experiences are shaped by the interplay of external sensory information, which becomes more robustly represented as the intensity of the experience increases, and internal cognitive processes, which provide the context and meaning for these experiences^43^. Ketamine’s effects on resting dynamics may therefore enhance access to critical internal information, fostering a neural state that supports more intense hedonic experiences, even when identical integrative processes are triggered by identical incoming inputs. This, however, suggests a complex three-way interaction: while greater information integration within the hedonic cluster is generally linked to stronger subjective experiences, the strength of this relationship may depend on ketamine’s effects on baseline neural dynamics.

To test this, we extracted averaged Φ^WMS^ values for each music trial (hedonic cluster) and baseline segment (baseline cluster) using edge masks derived from their respective NBS cluster results. A linear mixed model revealed the predicted significant three-way interaction between acute hedonic integration, shifts in baseline integration, and treatment (*β*_stand_ = 0.17, SE = 0.05, t(1251.29) = 3.14, p < 0.01; Table S7), supporting the idea that ketamine’s effects on baseline dynamics modulate the relationship between integration and subjective experience. In the placebo condition, the relationship between greater synergistic integration within the hedonic cluster and stronger hedonic experiences was unaffected by changes in the baseline cluster (Figure 2B). However, under ketamine, this relationship was significantly modulated: the ketamine-induced shift toward redundancy enhanced subjective experiences compared to placebo, even with similar levels of integration within the hedonic cluster. Figure 2C illustrates this effect more clearly, further showing that as subjective experiences intensify, the hedonic cluster shifts from redundancy-dominated to synergy-dominated activity.

### Opposite patterns of synergy and redundancy shape ketamine-induced effects on the hedonic experience

In general, changes in Φ^WMS^ (e.g., shifts toward synergy) can arise through three mechanisms: (i) increased synergistic processing (higher Φ^R^), (ii) reduced redundant processing (lower rtr), or (iii) a combination of both. To better understand the elementary dynamics driving these observed cluster effects, we used Integrated Information Decomposition^44^ (ΦID) to decompose Φ^WMS^ into its two components: Φ^R^ (synergistic and transfer processes) and rtr (redundant processes).

Using the hedonic NBS cluster edge masks, we extracted and averaged edge values for each trial, separating Φ^WMS^-increasing and Φ^WMS^-decreasing edges. This resulted in four metrics per trial: Φ^R+^ and rtr^+^ (the decomposition of the Φ^WMS^-increasing edges), Φ^R-^ and rtr^-^ (the decomposition of the Φ^WMS^-decreasing edges). We then fitted a linear mixed-effects model with all four decomposed dynamics per trial (Φ^R+^, rtr^+^, Φ^R-^, and rtr^-^) predicting the intensity of the hedonic experience and conducted a dominance analysis (DA) to assess the relative importance of these metrics in predicting changes in hedonic intensity^45^. The linear model shows that all four predictors significantly influence the intensity of the phenomenological experience, highlighting that hedonic experiences arise from a dynamic interplay between redundancy and synergy, rather than being driven solely by synergy (see Table S8). Focusing on the results of the DA reveals that changes in redundancy (rtr) were far more predictive of the subjective intensity of the hedonic experience than changes in synergy (Φ^R^) (see Table S9 and Figure 2D). This was particularly evident in the Φ^R+^ and rtr^+^ dynamics, for which decreases in redundancy were more strongly associated with changes in subjective hedonic experiences than increases in synergy. Thus, while both synergy and redundancy contribute to the emergence of hedonic experiences, the DA underscores that changes in redundancy play a considerably greater role in shaping these experiences.

Using a similar method, we investigated the dynamics underlying the nearly uniform decrease in Φ^WMS^ observed at rest after ketamine administration, which amplified the subjective hedonic experience. In this case, we fitted a logistic regression model with the four metrics (Φ^R+^, rtr^+^, Φ^R-^, and rtr^-^) to predict treatment. The model revealed that all four dynamics significantly predicted treatment (see Table S10). The DA highlighted that the Φ^R−^ and rtr^−^ metrics – associated with the majority of all significantly shifting edges after ketamine administration – jointly contributed to the overall shift toward redundancy (see Table S11 and Figure 2E). Both decreases in synergy and increases in redundancy were consistently highly predictive of treatment condition across all model subsets. A similar but weaker pattern was observed for the decomposed dynamics associated with increased Φ^WMS^ edges after ketamine.

### The dynamic neural processing signature underlying peak hedonic experiences

Next, we investigated the temporal evolution of information dynamics underlying hedonic experiences, hypothesizing a universal structure (across treatment) driven by one of two patterns: (i) remaining in or frequently transitioning to Φ^WMS+^ states, or (ii) alternating between Φ^WMS+^ and Φ^WMS-^ states, allowing the brain to flexibly integrate (Φ^WMS+^) and share redundant (Φ^WMS-^) information to support hedonic experiences. To test this, we concatenated run-wise Φ^WMS^ parcel-by-parcel time courses across participants and clustered them using a Gaussian mixture model. The optimal model identified 9 clusters: 3 integrative states (exclusively Φ^WMS+^), 3 redundant states (exclusively Φ^WMS-^), and 3 intermixed states (a mix of Φ^WMS+^ and Φ^WMS-^) (Figure 3A). From these brain state time series, we derived trial-wise transition matrices to quantify how often the brain transitioned between states. Finally, to explore the relationship between dynamic state switching and subjective experiences (that is, the moving experience, chills, and perceived valence), we utilized sparse canonical correlation analysis (CCA).

**Figure 3.**
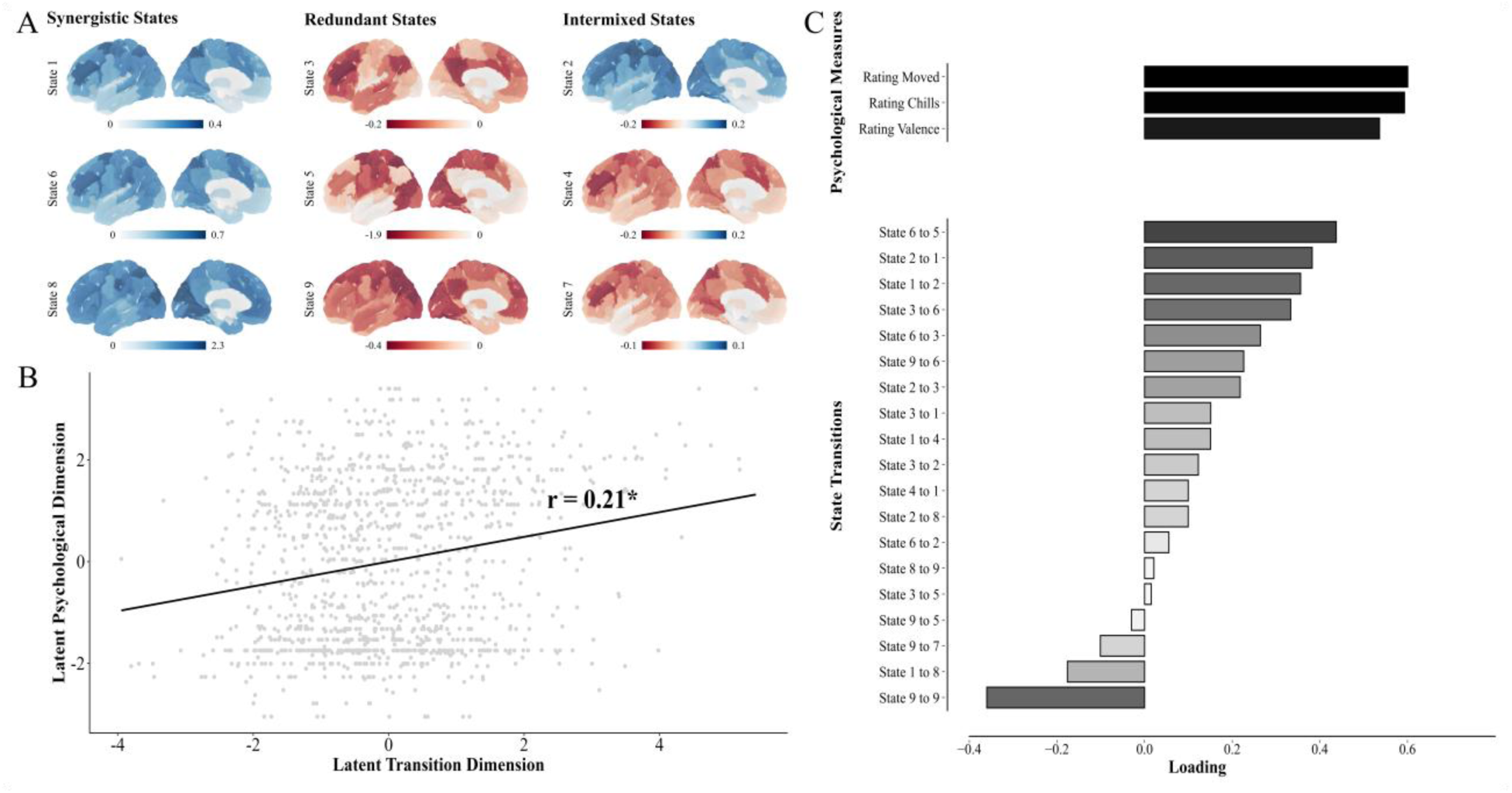
Hedonic experiences emerge from dynamic shifts between synergy and redundancy. (A) shows the clustering solution derived from the Gaussian Mixture Model applied to the recorded neural information dynamics across all participants (n = 32). Among the tested models (7 to 15 clusters), the optimal solution, based on the lowest BIC, was a 9-cluster model, revealing three distinct types of states: states characterized exclusively by increased synergy (left), those with increased redundancy (middle), and intermixed states (right). (B) presents the correlation between hedonic measures and state transitions within the recovered CCA latent space. (C) displays the canonical loadings from the Canonical Correlation Analysis (CCA) linking hedonic (psychological) measures and neural state transitions observed during the music trials. * indicates significance at p < 0.01.

The analysis revealed a meaningful shared structure between the two data sets (*r* = 0.21, *p* < 0.01, see Figure 3B). On the behavioral side, all three measures (moving, chills, and valence) loaded positively onto the latent dimension, reflecting a unified positive hedonic experience (Figure 3C). On the neural side, the regularized CCA reduced 81 transition features to 19 non-zero loadings (Figure 3C). The strongest negative loading corresponded to remaining in state 9, a globally redundant state, while all other self-transitions were zeroed out, suggesting that sustained occupancy of any single state – especially a redundant one – is not conducive to the emergence of peak hedonic experiences. Instead, dynamic switching appears critical. The strongest positive loading reflected transitions from state 6 (integrative) to state 5 (redundant), highlighting a sequence where integration is followed by stabilization through redundancy. Other key transitions, such as between state 3 (redundant) and state 6 (integrative), further emphasize the importance of the dynamic interplay between integration and redundancy in shaping hedonic experiences.

### The neurobiological mechanisms underlying changes in information integration

To elucidate the neurobiological mechanisms underlying the main interaction between ketamine-induced reductions in information integration at rest and the modulation of information integration during hedonic experiences, we examined the brain’s neurotransmitter systems. Specifically, we assessed whether spatial patterns of neurotransmitter systems align with the dominant integration changes identified in the NBS analysis. As distinct neurotransmitter systems might differently contribute to both decreases and increases in information integration, we only focused on the dominant shift of information integration in each identified NBS cluster. For the hedonic cluster, where Φ^WMS+^ edges dominated, we averaged the positive NBS edge weights for each node to create region-wise maps reflecting shifts toward synergy (see Figure 4). Conversely, for the baseline cluster, which was primarily driven by Φ^WMS-^ edges, we averaged the negative edge weights for each node to capture region-wise shifts toward redundancy. These node-level maps, representing the dominant integration changes within each cluster, were compared to neurotransmitter and transporter density maps^46^, including serotonin (5-HT1a, 5-HT1b, 5-HT2a, 5-HT4, 5-HT6, and 5-HTT), dopamine (D1, D2, and DAT), acetylcholine (M1, VAChT, and α4β2), glutamate (mGluR5 and NMDA), GABA (GABA), histamine (H3), cannabinoid (CB1), opioid (MOR), and norepinephrine transporter (NET). In addition, we also conducted an exploratory analysis to identify neurotransmitter systems that might underlie the ketamine-induced effects during music perception which is reported in the supplementary material.

**Figure 4.**
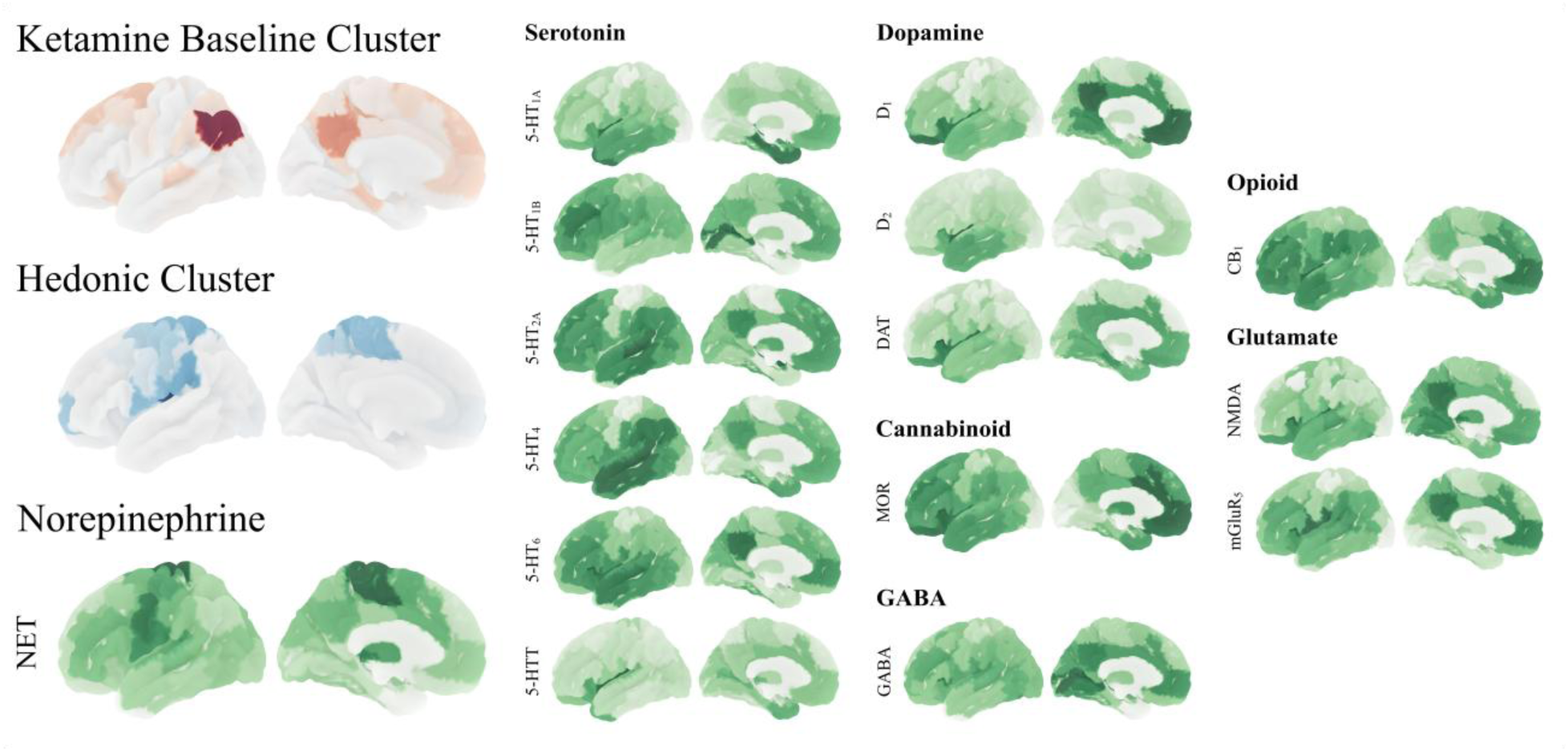
Synergistic hedonic integration is associated with norepinephrine transporter receptor density. Ketamine baseline cluster activity (in red) is shown by summing only the edges that increase in redundancy (Φ^WMS-^). Hedonic cluster activity (in blue) is shown by summing only the edges that present a significant increase in synergistic interactions (Φ^WMS+^). Neurotransmitter receptor and transporter density maps are depicted in green. For visual purposes, the significantly associated norepinephrine transporter density map is displayed alongside the hedonic cluster.

Our analysis revealed a significant association between NET density and Φ^WMS^ increases in the hedonic cluster (r = 0.53, *p* < 0.01), which survived Bonferroni correction for multiple comparisons (*p*_corrected_ = 0.02; see Figure 4). While for the ketamine-induced Φ^WMS^ reductions in the baseline cluster, we found no significant correlations with any receptor map.

## Discussion

While ketamine’s therapeutic effects are well-documented, the mechanisms underlying its pro-hedonic effects remain poorly understood. To address this gap, we developed a novel paradigm using self-selected music as a dynamic and emotionally rich stimulus to elicit a range of hedonic responses and applied information theory to characterize the resulting shifts in multivariate neural information integration. This approach allowed us to track how hedonic experiences unfold over time and how they are modulated by ketamine, providing a more nuanced understanding of both the hedonic experience and ketamine’s role in shaping it.

Our behavioral findings in healthy participants extend previous clinical research by showing that ketamine enhances all facets of the hedonic experience – ranging from deep emotional engagement to strong physiological responses – regardless of the initial valence of the stimulus. This universal amplification, even in response to mundane stimuli (e.g., neutral music), may help explain ketamine’s effectiveness in alleviating severe anhedonia in clinical populations that usually struggle to experience pleasure in their daily lives.

In the absence of external engagement, the brain’s global organization (as measured by O-information) is predominantly characterized by redundancy confirming findings from previous studies^47,48^. Yet, engagement with a complex stimulus shifts this organization toward greater synergy, reducing redundancy. This shift becomes even more pronounced during intense music-evoked hedonic experiences, suggesting that hedonic processing relies on the synergistic interaction of many widely distributed cortical and subcortical areas. Reminiscent of the brain’s underlying architecture for hedonic processes, these interactions are potent enough to reshape global information dynamics while preserving an overall redundant structure, reflecting a nuanced reconfiguration rather than a complete transformation of global neural dynamics.

So, are hedonic experiences the sole product of increases in synergistic processing, as initially hypothesized? Our findings suggest otherwise. Implementing Φ^WMS^ to analyze the temporal and spatial coevolution of more fine-grained information dynamics, we found that, in general, the pleasure response emerges from a localized balance between synergy and redundancy within a distinct hedonic cluster. Yet, as the hedonic experience intensifies, the overall balance does shift toward a synergy-dominated state, with the insular cortex emerging as a central hub, consistent with its established role in sensory integration^49^. However, primary auditory areas exhibit a different pattern, showing increased redundant coevolution with the insula, the thalamus, and the fusiform gyrus, a multisensory integration region involved in auditory emotion evaluation^50–52^. The observed redundancy ensures that auditory signals – the critical incoming information for the subsequently emerging hedonic experiences – are robustly transmitted to key relay and integration hubs. This pattern suggests that the brain prioritizes redundancy to safeguard the transmission of behaviorally critical information, making the information less error-prone and less susceptible to disruption for reliable and stable downstream processing. This interpretation aligns with neurobiological evidence in humans and animals, which underscores the role of redundancy in stabilizing essential sensory information^48,53–55^ and enhancing subsequent behavioral responses^21,22,56,57^. For instance, studies in animals have shown that more redundant encoding of crucial sensory information improves subsequent performance in decision-making tasks^56^, despite the higher energy cost of maintaining redundant neural representations. Our decomposition analysis further supports this notion, revealing that peak hedonic experiences are most robustly predicted by changes in redundant information processing (as measured by rtr). Considering the substantial energetic costs of sustaining a highly redundant system, minimizing unnecessary redundancy in large parts of the network is a logical means of reallocating resources more efficiently toward retaining crucial information that is indispensable for subsequent processing. This strategy is most evident in the synergy-increasing edges, which prominently reflect reductions in redundancy between the insula and primary sensory and motor cortices, with the notable exception of the auditory cortex. In addition, the topology of these changes in higher-order dynamics was associated with the distribution of norepinephrine transporter receptors, offering a neurobiologically plausible link between the emergence of a hedonic experience and norepinephrine^58^, a well-known neurotransmitter involved in modulating arousal, motivation, and euphoria, and dopamine^59^, an important neurotransmitter of the reward system While changes in redundancy are clearly crucial for the emergence of hedonic experiences, the interplay with synergy is equally important^54,60^, as demonstrated by our computational model. Peak hedonic experiences do not arise from prolonged occupancy of either redundancy or synergy-dominated states, but from dynamic transitions between the two. Synergy-dominated states enable efficient higher-order processing of complex information, while redundancy-dominated states stabilize and maintain these representations over time. This continuous alternation between synergistic integration and redundant stability may represent a general neural mechanism for balancing flexibility and reliability in ongoing experiences: redundancy ensures stable and reliable information access, while synergy allows the system to encode more information than the sum of its individual parts.

How does ketamine now facilitate its pro-hedonic effects? Various theories propose mechanisms such as changes in neurotransmitter systems^61^, relaxation of top-down control^62^, or neuroplasticity through synaptogenesis^63^, but they all converge on a common principle: ketamine must fundamentally alter neural information dynamics to create a state more conducive to the emergence of hedonic experiences. The key question, then, is how these effects are facilitated over the therapeutic timeframe. Are they driven by radical, global shifts in neural dynamics, or by more targeted, localized changes in information processing? While previous studies have shown that ketamine can indeed induce powerful global changes in information dynamics^64^, our results indicate that these effects are limited to the acute phase, and thus, are unlikely to underlie its long-term pro-hedonic effects. Although ketamine showed lingering sub-acute effects on localized information dynamics, it did not enhance local information integration directly linked to the strength of the subjective experience during stimulus processing, as no interaction effect between intensity of the experience and treatment was found. Instead, our findings point to a more foundational mechanism: ketamine induces subtle, subacute shifts in resting dynamics, (almost) uniformly shifting information dynamics toward greater redundancy across its network.

Interestingly, the greater this shift toward redundancy, the stronger hedonic experiences became in response to the same level of hedonic integration. This shift appeared to result from both increases in redundancy but also decreases in synergy, with both equally predictive of the intensity of the phenomenological experience. Just as encoding critical *external* information more redundantly benefits subsequent (hedonic) processes, ketamine-induced redundant dynamics at rest may similarly enhance the brain’s ability to reliably relay and represent critical *internal* information, which in turn supports the emergence of stronger hedonic experiences. This aligns with the idea that hedonic experiences are shaped by both the properties of external stimulation and internal cognitive processes. As basic research has shown, the intensity of a hedonic experience significantly increases when the stimulus is embedded in a rich context with a wide range of knowledge-related or personal associations^65,66^. Robust and widespread access to this internal information is therefore crucial. Ketamine’s long-term pro-hedonic effects likely stem from its ability to enhance the reliable representation of (critical) internal information, creating a neural environment optimized for subsequent hedonic processing.

As it is the case for all studies, our work is not without limitations. First, instead of using a classical resting-state scan that is recorded for multiple minutes and detached from the task, we employed interleaved resting periods throughout the task. Although unconventional, these interwoven periods, with eyes closed and no task instructions, were necessary to better link changes in default information dynamics to increased subjective experiences during task engagement. Notably, there is already some prior evidence showing that ketamine increases redundancy in resting-state dynamics^67,68^, providing additional support for our hypothesis. Second, while the use of self-selected music may have introduced variability across participants, it was crucial for eliciting strong hedonic experiences tailored to the individual. Nonetheless, our within-subject design, combined with linear mixed models, ensured robust and reliable results despite this variability. Finally, our conjecture that ketamine’s pro-hedonic effects arise from its impact on intrinsic dynamics at rest, enhancing access to internal information, can be easily tested in future studies by examining whether these effects diminish as redundancy returns to baseline and whether this improved access to internal information enhances performance in other tasks as well for which widespread information access is equally essential.

Collectively, our data reveals that hedonic experiences arise from a nuanced interplay between synergistic and redundant information processes. By increasing a redundancy-dominated organization at rest, ketamine facilitates an optimized infrastructure by inducing stable and widespread communication of information. Once external information enters the system, resources can be flexibly reallocated to process critical information from anywhere in the brain, stabilizing higher-order integrative dynamics^69^ essential for the hedonic experience, and thus, maximizing the behavioral readout. As a result, even modest increases in information integration during music perception produce disproportionately strong hedonic experiences. Thus, ketamine’s pro-hedonic effects likely stem from its ability to boost the brain’s capacity to reliably represent critical internal information, fostering a neural environment more conducive to the emergence of hedonic experiences.

## Acknowledgements

The authors extend their heartfelt gratitude to the students, medical professionals, and technicians whose contributions were essential to the successful implementation of this study. This work would not have been possible without the collaborative efforts of the Vienna Cognitive Science Hub and the Advanced Neuroimaging Lab.

## Methods

### Study design

This randomized, single-blind, cross-over, placebo-controlled study enrolled 40 healthy participants between the age of 18 and 55. Physical health was assessed through medical examinations including electrocardiogram, blood, urine, pregnancy, and drug tests, while mental health was evaluated using the Structured Clinical Interview^70^. Exclusion criteria included history of neurological and psychiatric disorders, any current pharmacological treatment, pregnancy, breastfeeding, current or former substance abuse and prior experience with ketamine (in context of recreational or therapeutic use).

Participants underwent two functional magnetic resonance imaging (fMRI) sessions, separated by at least two weeks to avoid cross-over effects (see Figure 1B). Four hours prior to each scanning session, participants received either a saline solution (0.9% NaCl) or 0.5mg/kg of racemic ketamine diluted in 0.9% NaCl, administered over 40 minutes. Participants performed a hedonic music task during each scan session (see Hedonic Task). Two scanner malfunctions occurred after drug administration was finished, preventing data collection of two participants. These participants were excluded from the study, resulting in a dataset of *N* = 38 participants. The study was approved by the ethics committee of the Medical University of Vienna (EK 1572/2021).

### Hedonic task

Prior to study inclusion, participants provided 10 self-selected highly positive (moving) songs and 10 self-selected neutral songs, indicating the exact moment in which the hedonic experience peaked during each positive song. Self-selection was used to account for individual variability in emotional responses to music^71^, as the focus was on eliciting robust hedonic experiences rather than controlling for music type. Note, neutral songs were included to determine whether ketamine broadly amplifies hedonic responses, regardless of their initial valence, or whether it enhances the hedonic experience only for stimuli already perceived as enjoyable. Participants were given detailed definitions of the concept of moving and chills to minimize semantic misunderstandings. Using ffmpeg^72^, 70-second excerpts were created around the individually defined peak moment for positive songs, while random intervals were selected for neutral songs. All excerpts were sampled at 44,100 Hz, volume-normalized, and edited with 2-second fade-ins and fade-outs to ensure smooth transitions.

Before scanning, participants reviewed task instructions, passed comprehension checks, and completed a sound test using Mozart’s “Eine kleine Nachtmusik” to optimize music volume in relation to scanner noise. The hedonic task was divided into two runs, each consisting of three blocks alternating between neutral and positive songs, with the starting condition and block size sequence randomized across participants. Positive and neutral songs were randomly assigned to two blocks of three songs and one block of four songs each. Each music trial began with a one-second pictogram of a closed eye, signaling participants to close their eyes for the upcoming trial (see Figure 1A). After listening to each excerpt, participants had 12 seconds to rate their hedonic experience on three different dimensions: a) moving— how moving the experience was on a scale of 1 to 10; b) chills—whether a chill was induced and how intense the chill was on a scale of 1 (no chill at all) to 10 (utmost chilling experience); and valence—how unpleasurable/pleasurable the experience was on a scale of −5 to 5. To mitigate ceiling effects, they were instructed to compare their experiences to past events that were most moving, chill-inducing, or pleasurable/unpleasurable. While valence reflects the general positivity or negativity of the experience, the neuroaesthetic concept of moving captures deep emotional engagement, which may or may not lead to strong physiological responses such as chills^73,74^. Thirty-second baseline (resting) segments with eyes closed were included before and after each block.

### MRI data acquisition

Scan sessions were performed roughly 4 hours after drug administration on a 3 Tesla Siemens MAGNETOM Prisma scanner at the Department of Biomedical Imaging and Image-guided Therapy at the Medical University of Vienna. Structural T1-weighted MRI scans were acquired with a multi-echo MPRAGE sequence (TR = 2400ms, TE = 2.22ms, voxel size 0.8×0.8×0.8mm, anterior-posterior face encoding, field of view read: 256mm). Functional data acquisition used a 64-channel head coil with a multiband 4 sequence for the hedonic task (TR = 800ms, TE = 30ms, voxel size 3×3×3mm, posterior-anterior phase-encoding direction, field of view read: 210mm). To aid distortion correction, short sequences in reversed encoding directions were recorded before each task run.

### Behavioral analyses

To assess whether ketamine merely affects the hedonic tone (valence) or also influences deeper affective (moving) and physiological experiences (chills), three independent hierarchical ordinal regressions with a logit link function were conducted using the ordinal package^75^ in R (n = 38). Consistent with Bürkner & Vuorre’s^76^ recommendations, we evaluated potential models based on different combinations of regressors (treatment, initial judgement, order of administration, and sex) and hierarchical effects. Comparisons were made based on the Bayesian Information Criteria (BIC) calculated with the performance package^77^. The winning model included a regressor for treatment (ketamine vs placebo), a regressor for the initial judgement (indicating whether the song was originally selected as neutral or positive), and a random effect for each participant (subject), accounting for the nestedness of the data. Notably, the winning model did not contain an interaction term between treatment and initial judgement. We fit a separate model for each of the three hedonic dimensions: moving, chills, and valence (see Hedonic Task). All model assumptions were visually inspected using the performance package^77^. To account for multiple comparisons, a Bonferroni correction was applied, adjusting the alpha significance level to 0.016.

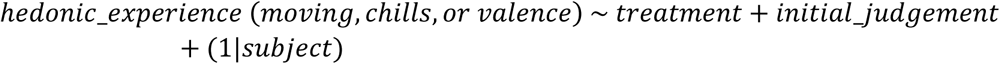

### MRI data preprocessing

Structural and functional imaging data were preprocessed using fMRIprep^78,79^ with default parameters. The structural preprocessing pipeline included intensity non-uniformity correction using N4BiasFieldCorrection^80^, brain tissue segmentation, and non-linear spatial normalization to MNI 2mm standard space using ANTS^81^. The MNI template was obtained through TemplateFlow^82^.

The functional preprocessing pipeline consisted of slice-timing correction, motion correction using MCFLIRT^83^, and B0 inhomogeneity correction using fieldmaps derived from runs with reversed encoding directions processed via TOPUP^84^. Functional data was co-registered to the T1-weighted image using bbregister^85^ and high-pass filtered with discrete cosine filters (128-second cut-off). Additional noise reduction was performed using aCompCor^86^. After high-pass filtering, principal components were calculated separately for white matter (WM) and cerebrospinal fluid (CSF) masks. The top five components from each mask, along with 24 motion parameters (six motion parameters, their quadratic terms, their first and second order derivatives), were regressed out using Nipype^87^.

Quality control criteria excluded datasets with a mean framewise displacement (FD) exceeding 0.2 mm, more than 20% outlier volumes, or a maximum FD greater than 5 mm^88^. Based on these thresholds, six participants were excluded resulting in a final sample size for the fMRI data analyses of 32 participants. Functional data was parcellated into 116 regions using an atlas combining both the cortical Schaefer100^33^ and the subcortical Tian16^34^ parcellations.

### O-Information analysis

The O-Information (also denoted as Ω) is a scalable measure to assess the interactions of a set of random variables^23^. Although it cannot disentangle separate contributions of synergy and redundancy, the O-Information captures the balance between the two: a system with Ω > 0 is characterized by information being shared across the system and is redundancy-dominated, and a system with Ω < 0 contains irreducible higher-order interactions and is synergy-dominated. Formally, Ω is derived from two generalized measures of mutual information: the total correlation (TC)^89^, which quantifies the collective constraints among a set of variables, and the dual total correlation (DTC)^90^, which assesses shared randomness across these variables. For a system of *n* random variables X^n^ = {X_1_,…, X_n_}, these measures are defined as:

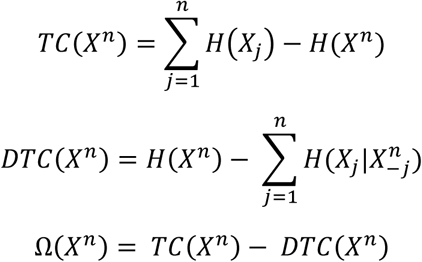

Here, H(X^n^) is the joint Shannon entropy of the *n*-plet, H(X_j_) is the entropy of the *j*-th variable, and H(X_j_|X^n^_-j_) is the conditional entropy of the *j*-th variable given the rest of the system.

To assess whether peak hedonic experiences modulate global neural information dynamics, we calculated O-Information for the 116-plet containing all brain regions simultaneously. Cleaned, parcellated BOLD data were concatenated across runs and participants and analyzed using the Gaussian copula solver from the JIDT toolbox^91^. This method provided localized O-Information values over time, capturing the balance between redundancy and synergy in neural activity. To account for the hemodynamic lag, O-Information values were extracted for each music trial and baseline segment by shifting the extraction window by 4 TRs. These values were then averaged within each extracted period to yield a single mean O-Information value for each music trial and baseline segment.

To explore the impact of music perception and subjective experiences on global neural processing, we fitted two linear mixed-effects models (n = 32) using the lme4 package^92^. The first model tested the effects of the regressors music perception (music vs baseline) and treatment (ketamine vs placebo) on O-Information, with participants included as a random effect. This model tested the implicit assumption that the engagement with a complex stimulus such as music would shift global neural dynamics toward greater synergy compared to rest.To account for multiple comparisons, a Bonferroni correction was applied, adjusting the alpha significance level to 0.025.

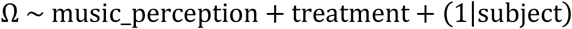

The second model examined how the strength of the hedonic experience – using the moving rating as our main measure, given its role as a deep positive emotional response – and treatment (ketamine vs placebo) influenced O-Information, with participants again included as a random effect to account for the nestedness of the data.

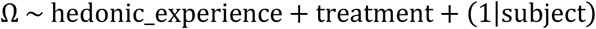

All model assumptions were visually inspected using the performance package^77^.

### Calculation and decomposition of Φ^WMS^ time courses

At its core, mutual information quantifies the interdependence of two variables X and Y. Specifically, mutual information I(X;Y) quantifies the amount of information that the knowledge of node X provides about node Y. Recently, mutual information was extended to systems of three or more variables using Partial Information Decomposition^93^. In this higher order case, PID decomposes the mutual information I(X,Y;Z) which the two source nodes X and Y provide in respect to a third target node Z into three distinct types of information. Information is considered *unique* (uni) if it is only provided by one of the sources but not the other, whereas *redundant* (red) information denotes information shared by both sources.

Lastly, *synergistic* (syn) information arises from the joint interaction of both nodes and cannot be reduced to any node individually.

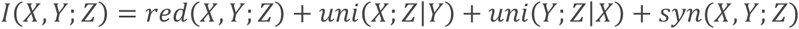

ΦID^27^ further extends PID by decomposing time-delayed mutual information (TDMI), both in the past (t_-1_) and in the future (t). For a pair of nodes, ΦID thus captures how information dynamics evolve over time, considering both the past and present state of the nodes I(X_t-1_,Y_t-1_;X_t_,Y_t_). In practice, this requires constructing a linear system of 15 equations with 16 unknowns, linking the standard Shannon mutual information to the redundant, unique, and synergistic components of TDMI. To be able to solve this set of equations, we utilized the widely used minimum mutual information^94^ (MMI) definition of redundancy:

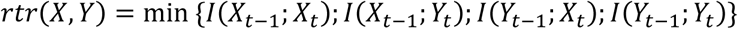

This temporal decomposition produces 16 distinct information atom time courses, capturing how information is transferred between nodes over time. For example, the atom rtr represents information that was shared by two sources in the past and remains shared in the future. Recent work^95^ has shown that *whole-minus-sum integrated information* (Φ^WMS^) can be expressed as the difference between a specific set of atoms

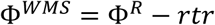

*Where:*

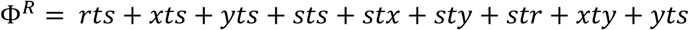

Here, we use the standard shorthand notation for the atoms introduced in prior work^95^, e.g. xts represents the atom describing how unique information from the source node X becomes synergistic over time.

For each run, preprocessed and parcellated BOLD data from all pairwise combinations of brain regions was fully decomposed using ΦID, yielding the time courses of all 16 atoms for each pair of regions. We then constructed Φ^R^ and Φ^WMS^ time courses by summing the relevant atoms. Notably, while Φ^R^ requires defining a redundancy function (here, MMI), Φ^WMS^ does not, as it is fully determined by the TDMI. However, we calculated it through the atoms for convenience. The analysis was performed using scripts provided by Luppi et al.^44^ and the Gaussian estimator from the JIDT toolbox^91^.

### Network-Based Statistic (NBS) analysis

To explore how the intensity of the hedonic experience and ketamine influence local neural information integration, we analyzed Φ^WMS^ time courses from music trial and baseline segments, adjusting for the hemodynamic lag by shifting the time series by three time points. Note, the time series was shifted by only 3 TRs because the Φ^WMS^ calculation inherently accounts for how information flows from one time step to the next, resulting in a time series that is already effectively shifted by 1 TR compared to the underlying BOLD signal. These time courses were averaged within each segment to create node-by-node Φ^WMS^ matrices for each condition.

We used a modified version of the Network-Based Statistic (NBS) algorithm^96^, based on linear mixed-effects models, which accounts for subject variability, unbalanced designs, and allows the simultaneous inclusion of both positive and negative edges within a single cluster. Briefly, NBS fits a linear model to each connection (edge) and identifies those exceeding a predefined statistical threshold. It then checks whether these significant edges form connected networks through shared nodes. The significance of each network cluster is determined using permutation testing, which corrects for multiple comparisons at the network level.

To examine how hedonic intensity and ketamine influence local information integration during music perception, we fitted a linear mixed-effects model to each connection with the Φ^WMS^ values of the music trial as the dependent variable, including regressors for treatment (ketamine vs placebo), the hedonic experience (the intensity of the moving experience), their interaction, and a random effect for each participant. Note, the cluster linked to the main effect of hedonic experience is termed the *hedonic cluster*, and the one linked to the main effect of treatment during music perception is referred to as the *ketamine trial cluster*.

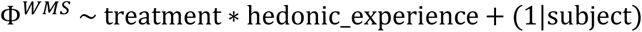

To investigate the effects of ketamine independent of music stimulation, we applied a separate linear mixed-effects model to the baseline (resting) segments. This model included a fixed effect for treatment (ketamine vs placebo) and a random effect for participants. The cluster associated with the main effect of treatment during the baseline segments is referred to as the *ketamine baseline cluster* or just *baseline cluster*.

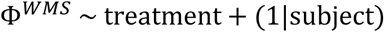

In our analysis (n = 32), we used 5,000 permutations and set the threshold to reflect a medium effect size (Cohen’s d = 0.5), corresponding to a t-value of 2.8. We used the intensity of each cluster, that is, the sum of t-statistics across connected edges, as the primary statistic for significance testing.

Importantly, the results remained consistent when using extent, that is, the number of edges in the cluster, supporting the robustness of our findings regardless of the specific NBS metric.

### Linking baseline and hedonic cluster activity to the intensity of the hedonic experience

We next considered whether ketamine’s pro-hedonic effects could arise from its ability to shift intrinsic dynamics at rest greater redundancy, thereby enhancing the brain’s capacity to reliably access key internal representations. Specifically, we examined whether persistent, ketamine-induced changes in baseline neural integration (ketamine baseline cluster) modulate the relationship between experience-evoked changes in information integration (hedonic cluster) and the phenomenological experience.

For each music trial, we extracted and averaged Φ^WMS^ edge values associated with the hedonic cluster, resulting in a single integration score per trial (avg_hedonic_cluster). Likewise, for each interleaved baseline segment, we extracted edge values from the ketamine baseline cluster and aggregated them into a single baseline integration score per run (avg_ketamine_baseline_cluster). This baseline score reflects persistent drug-induced changes in information integration at rest, independent of music stimulation.

To test whether the activity of these two clusters jointly predicts hedonic responses, we fitted a linear mixed-effects model with the reported hedonic experience (here, the intensity of the moving experience) as the dependent variable. The model (n = 32) included the average hedonic cluster activity per trial, the average change in baseline cluster activity, treatment (ketamine vs placebo), and their three-way interaction as fixed effects, with a random intercept per participant.

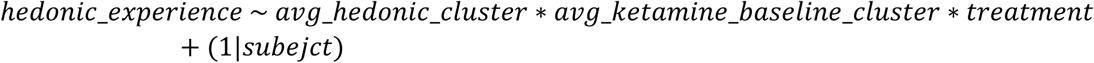

To assess statistical significance, we conducted 10,000 wild bootstrap iterations using the lmersampler^97^ package in R, which accounts for the hierarchical structure in our data.

### Dominance analysis (DA)

To investigate the underlying dynamics driving changes in Φ^WMS^, we performed a dominance analysis. First, we created separate edge masks for the increasing and decreasing Φ^WMS^ edges identified in the NBS analysis for both the hedonic cluster and the ketamine baseline cluster. We then extracted and averaged the edge values for each trial and baseline segment from the decomposed time series (Φ^R^ and rtr), resulting in four metrics per trial/segment: Φ^R+^ and rtr^+^ (representing the decomposition of Φ^WMS^-increasing edges) and Φ^R-^ and rtr^-^ (representing the decomposition of Φ^WMS^-decreasing edges).

Next, we fitted a linear mixed-effects model to predict the intensity of hedonic experiences (here, moving ratings) using these four trial-averaged dynamics, with a random intercept for each participant:

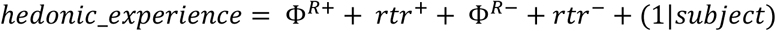

The model results were corrected using cluster-robust covariance matrices (CR1) via the sjPlot package^98^. To further assess the relative importance of each dynamic, we conducted a bootstrap dominance analysis with 1,000 iterations of the model parameters using the dominanceanalysis package in R^99^.

Similarly, we fitted a generalized linear model (GLM) with a logit link function to predict treatment assignment based on the four dynamics from the baseline segments and conducted a dominance analysis of the model parameters. An earlier version of this model included a random intercept for participants, but it was removed after determining that the random intercept was singular (close to zero) and unnecessary.

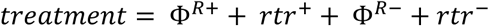

### Association with neurotransmitter density maps

To investigate whether spatial patterns of neurotransmitter receptor and transporter densities align with the ketamine-induced reductions in information integration at rest and the modulation of information integration during hedonic experiences, we conducted a spatial correlation analysis. Since different neurotransmitter systems may uniquely affect increases and decreases in information integration, we concentrated on the dominant shift in each identified NBS cluster. As both NBS-identified clusters contained a mix of positively and negatively weighted edges, we first computed the net effect within each cluster by subtracting the average weight of negative edges from that of positive edges. This allowed us to determine the dominant direction of change in the cluster. The hedonic cluster during music perception was dominated by Φ^WMS+^ edges, whereas the ketamine cluster during the baseline segments was almost exclusively comprised of Φ^WMS-^ edges. As expected, in both cases, the number of edges in the dominant direction also exceeded those in the opposite direction. We did this based on the reasoning that different neurotransmitter systems may underlie increases versus decreases in Φ^WMS^, thus focusing our analysis on the dominant direction of each cluster. To create node-level effects for these two observed changes in information dynamics, we collapsed the observed edge dynamics from the NBS analyses by averaging the weights (t-values) of each significantly identified edge per node. These integration maps were compared against PET-derived neurotransmitter and transporter density maps taken from Hansen et al.^100^, which were parcellated using the combined Schaefer100^33^ and Tian16^34^ parcellation scheme, consistent with the rest of our analysis. Tested density distributions included maps for serotonin (5-HT1a^101^, 5-HT1b^101,102^, 5-HT2a^103^, 5-HT4^103^, 5-HT6^104^, and 5-HTT^101,103^), dopamine (D1^105^, D2^106,107^, and DAT^108^), acetylcholine (M1^109^, VAChT^110,111^, and α4β2^112^), glutamate (mGluR5^100,113^ and NMDA^100^), gamma-aminobutyric acid (GABAa^114^), histamine (H3^102^), cannabinoid (CB1^115^), opioid (MOR^116,117^), and norepinephrine transporter (NET^118,119^).

To test the significance of the similarity between neural integration maps and receptor maps, while accounting for the innate spatial autocorrelation of receptors, we created receptor surrogate maps using a variogram-based approach^120^. Compared to traditional spin tests, this method is suitable for whole-brain and particularly parcellated data and has been shown to better preserve the intrinsic spatial autocorrelation of brain maps. Briefly, for each receptor or transporter map, a variogram was computed to quantify the relationship between pairwise variance in parcel density values as a function of their spatial distance, capturing the map’s inherent spatial autocorrelation structure. The original map is then randomly shuffled, destroying this structure, and subsequently smoothed using a kernel, which reintroduces a new spatial autocorrelation structure. An iterative optimization process is then used to identify the smoothing kernel that minimizes the deviation between the smoothed surrogate’s variogram and the original map’s variogram (quantified as the sum of squared errors). The best-fitting kernel is then used to create one surrogate map with shuffled values, yet the same spatial autocorrelation as the original map. We generated 10,000 surrogate maps per receptor/transporter and computed their Spearman’s rank correlations with the integration maps derived from the ketamine baseline cluster and the hedonic cluster. Results were corrected for multiple comparisons using the Bonferroni correction method. Additionally, an exploratory analysis was performed by separately examining the node-averaged edge weights of the positively and negatively weighted connections within the NBS cluster linked to the ketamine effect during music perception (see Supplemental Material).

### Canonical correlation analysis (CCA)

To investigate whether dynamic changes between brain state-like integration patterns are linked to the emergence of hedonic experiences, we first concatenated run-wise Φ^WMS^ time courses across participants and clustered the resulting data using a Gaussian Mixture Model^121^. We evaluated models with 7 to 15 clusters and selected the optimal number based on BIC. The best-fitting solution consisted of 9 clusters, each representing a recurring pattern of neural information dynamics.

Next, we computed state transition probabilities for each music trial, that is, the likelihood of transitioning from one brain state to another (or remaining in the same state), resulting in 81 transition probabilities per trial. To explore whether these transition dynamics were meaningfully related to participants’ reported hedonic experiences, we employed Canonical Correlation Analysis (CCA). CCA is a multivariate statistical technique used to identify shared variance between two sets of variables – in this case, brain-derived state transition probabilities and hedonic experiences (all three behavioral ratings). It does so by finding linear combinations (called canonical variates) within each dataset that are maximally correlated with one another. Formally, CCA identifies weight vectors that maximize the correlation between these linear combinations, i.e., between the projections of the brain data and behavioral data into a shared latent space. In essence, it reveals a latent dimension that best explains the covariation between the neural and behavioral data.

To determine the appropriate number of canonical dimensions to retain, we first applied Principal Component Analysis and used the elbow method as a visual heuristic. This analysis revealed a single meaningful dimension, explaining 10% of the neural variance and 85% of the behavioral variance. Based on this, we limited our CCA to a single latent canonical dimension. Given the high dimensionality of the neural dataset and the potential for multicollinearity, we implemented a regularized CCA using the PMA package in R^122^. Regularization was applied exclusively to the standardized neural data to improve model interpretability. We iteratively tuned the L1 sparsity parameter from 0.1 to 1.0 in increments of 0.05, using the internal permutation-based tuning function in PMA. Briefly, this procedure permutes the neural dataset 25 times, runs sparse CCA for each permutation and the original data, and then calculates the correlation between the resulting latent variables. These correlations are Fisher-Z-transformed to approximate a normal distribution, and a Z-statistic is computed by comparing the observed correlation against the distribution of permuted results. The tuning parameter with the highest Z-statistic was selected as the optimal level of sparsity. Since we were interested in identifying whether a unique pattern of state transitions is linked to hedonic experiences independently of treatment, we assessed the significance of the CCA results by randomly shuffling behavioral ratings within participants 10,000 times. Specifically, we reassigned complete trial-level ratings to different neural transition matrices. This approach preserved the internal structure of the data while disrupting the correspondence between neural and behavioral variables (as well as their alignment across treatment), thereby generating a null distribution of canonical correlations against which the observed correlation could be compared.

