## Supplemental Material for "Hedonic experiences emerge from an orchestrated balance of synergistic and redundant information processing"

| Valence | | | | | |
|---|---|---|---|---|---|
| *Predictors* | *Odds Ratios* | *SE* | *CI (95%)* | *z-statistic* | *p-Value* |
| -5\|-4 | 0.01 | 0.00 | 0.01 – 0.02 | -12.77 | **<0.001** |
| -4\|-3 | 0.03 | 0.01 | 0.01 – 0.04 | -13.49 | **<0.001** |
| -3\|-2 | 0.05 | 0.01 | 0.03 – 0.08 | -12.75 | **<0.001** |
| -2\|-1 | 0.14 | 0.03 | 0.10 – 0.21 | -9.57 | **<0.001** |
| -1\|0 | 0.41 | 0.08 | 0.28 – 0.60 | -4.63 | **<0.001** |
| 0\|1 | 3.56 | 0.69 | 2.44 – 5.20 | 6.59 | **<0.001** |
| 1\|2 | 18.16 | 3.88 | 11.94 – 27.62 | 13.56 | **<0.001** |
| 2\|3 | 92.99 | 21.72 | 58.84 – 146.98 | 19.41 | **<0.001** |
| 3\|4 | 368.66 | 91.14 | 227.09 – 598.50 | 23.91 | **<0.001** |
| 4\|5 | 1547.13 | 416.24 | 913.11 – 2621.39 | 27.30 | **<0.001** |
| Drug condition [Ketamine] | 1.36 | 0.13 | 1.13 – 1.63 | 3.24 | **0.001** |
| Initial Judgment [Positive] | 116.08 | 18.61 | 84.77 – 158.94 | 29.65 | **<0.001** |
| **Random Effects** | | | | | |
| $\sigma^2$ | 3.29 | | | | |
| $\tau_{00\ ID}$ | 1.03 | | | | |
| ICC | 0.24 | | | | |
| $N_{ID}$ | 38 | | | | |
| Observations | 1501 | | | | |

Table S1: Ordinal Regression Results for Valence Ratings
This table presents the results of the cumulative ordinal regression analysis on valence ratings using a logit link function. The model includes drug condition (placebo, ketamine) and initial judgment (neutral, positive) as predictors, with a random intercept for each participant to account for repeated measures. Exponentiated parameter estimates (Odds Ratios) along with their 95% confidence intervals (CI) and standard error (SE) are reported.

| Moving | | | | | |
|---|---|---|---|---|---|
| *Predictors* | *Odds Ratios* | *SE* | *CI (95%)* | *z-statistic* | *p-Value* |
| 1\|2 | 0.45 | 0.08 | 0.32 – 0.64 | -4.48 | **<0.001** |
| 2\|3 | 1.14 | 0.20 | 0.80 – 1.60 | 0.72 | 0.472 |
| 3\|4 | 2.78 | 0.50 | 1.95 – 3.95 | 5.69 | **<0.001** |
| 4\|5 | 5.45 | 1.01 | 3.79 – 7.84 | 9.14 | **<0.001** |
| 5\|6 | 10.29 | 2.00 | 7.03 – 15.05 | 12.00 | **<0.001** |
| 6\|7 | 44.21 | 9.75 | 28.69 – 68.13 | 17.18 | **<0.001** |
| 7\|8 | 333.51 | 81.47 | 206.62 – 538.32 | 23.78 | **<0.001** |

| | | | | | |
|---|---|---|---|---|---|
| 8\|9 | 1548.58 | 404.37 | 928.25 – 2583.46 | 28.13 | **<0.001** |
| 9\|10 | 7022.94 | 2115.09 | 3891.91 – 12672.85 | 29.41 | **<0.001** |
| Drug condition [Ketamine] | 1.31 | 0.12 | 1.09 – 1.58 | 2.88 | **0.004** |
| Initial Judgment [Positive] | 219.68 | 39.28 | 154.74 – 311.87 | 30.16 | **<0.001** |
| **Random Effects** | | | | | |
| $\sigma^2$ | 3.29 | | | | |
| $\tau_{00\ ID}$ | 0.87 | | | | |
| ICC | 0.21 | | | | |
| $N_{ID}$ | 38 | | | | |
| Observations | 1488 | | | | |

Table S2: Ordinal Regression Results for Moving Ratings
This table presents the results of the cumulative ordinal regression analysis on moving ratings using a logit link function. The model includes drug condition (placebo, ketamine) and initial judgment (neutral, positive) as predictors, with a random intercept for each participant to account for repeated measures. Exponentiated parameter estimates (Odds Ratios) along with their 95% confidence intervals (CI) and standard error (SE) are reported.

| | **Chills** | | | | |
|---|---|---|---|---|---|
| *Predictors* | *Odds Ratios* | *SE* | *CI (95%)* | *z-statistic* | *p-Value* |
| 1\|2 | 4.09 | 1.13 | 2.37 – 7.03 | 5.08 | **<0.001** |
| 2\|3 | 8.69 | 2.44 | 5.01 – 15.08 | 7.69 | **<0.001** |
| 3\|4 | 17.51 | 5.03 | 9.97 – 30.75 | 9.96 | **<0.001** |
| 4\|5 | 34.76 | 10.26 | 19.48 – 62.00 | 12.01 | **<0.001** |
| 5\|6 | 66.47 | 20.17 | 36.67 – 120.49 | 13.83 | **<0.001** |
| 6\|7 | 274.02 | 87.45 | 146.60 – 512.17 | 17.59 | **<0.001** |
| 7\|8 | 1123.91 | 374.64 | 584.78 – 2160.06 | 21.07 | **<0.001** |
| 8\|9 | 3452.90 | 1213.63 | 1733.81 – 6876.46 | 23.18 | **<0.001** |
| 9\|10 | 13886.20 | 5725.81 | 6188.72 – 31157.74 | 23.13 | **<0.001** |
| Drug condition [Ketamine] | 1.60 | 0.17 | 1.31 – 1.96 | 4.52 | **<0.001** |
| Initial Judgment [Positive] | 114.32 | 18.62 | 83.07 – 157.33 | 29.09 | **<0.001** |
| **Random Effects** | | | | | |
| $\sigma^2$ | 3.29 | | | | |
| $\tau_{00\ ID}$ | 2.46 | | | | |
| ICC | 0.43 | | | | |
| $N_{ID}$ | 38 | | | | |
| Observations | 1499 | | | | |

Table S3: Ordinal Regression Results for Chills Ratings
This table presents the results of the cumulative ordinal regression analysis on chills ratings using a logit link function. The model includes drug condition (placebo, ketamine) and initial judgment (neutral, positive) as predictors, with a random intercept for each participant to account for repeated measures. Exponentiated parameter estimates (Odds Ratios) along with their 95% confidence intervals (CI) are reported.

| | O-Information Music vs Rest | | | | | | | |
|---|---|---|---|---|---|---|---|---|
| *Predictors* | *Estimates* | *SE* | *std. Beta* | *std. SE* | *CI (95%)* | *std. CI (95%)* | *t-statistic* | *p-Value* |
| Intercept | 32.39 | 0.67 | 0.09 | 0.10 | 31.08 – 33.71 | -0.11 – 0.29 | 48.21 | **<0.001** |
| Music or Rest [Music] | -0.73 | 0.30 | -0.11 | 0.04 | -1.32 – -0.15 | -0.20 – -0.02 | -2.46 | **0.014** |
| Drug Condition [Placebo] | -0.13 | 0.27 | -0.02 | 0.04 | -0.66 – 0.39 | -0.10 – 0.06 | -0.49 | 0.623 |
| **Random Effects** | | | | | | | | |
| σ2 | 32.18 | | | | | | | |
| τ00 subject | 11.84 | | | | | | | |
| ICC | 0.27 | | | | | | | |
| N subject | 32 | | | | | | | |
| Observations | 1782 | | | | | | | |

Table S4: Linear mixed-effects model for O-Information across rest and music perception
The model includes current state (rest, music perception) and drug condition (placebo, ketamine) as predictors, with a random intercept for each participant to account for repeated measures. Unstandardized coefficients (Estimates) and standardized beta estimates (std. Beta), along with their 95% confidence intervals (CI) and standard error (SE), are reported.

| | O-Information Rest Segments | | | | | | | |
|---|---|---|---|---|---|---|---|---|
| *Predictors* | *Estimates* | *SE* | *std. Beta* | *std. SE* | *CI (95%)* | *std. CI (95%)* | *t-statistic* | *p-Value* |
| Intercept | 32.25 | 0.80 | -0.01 | 0.10 | 30.68 – 33.82 | -0.20 – 0.18 | 40.36 | **<0.001** |
| Drug Condition [Placebo] | 0.13 | 0.67 | 0.02 | 0.08 | -1.18 – 1.45 | -0.14 – 0.17 | 0.20 | 0.843 |
| **Random Effects** | | | | | | | | |
| σ2 | 56.79 | | | | | | | |
| τ00 subject | 13.31 | | | | | | | |
| ICC | 0.19 | | | | | | | |
| N subject | 32 | | | | | | | |
| Observations | 508 | | | | | | | |

Table S5: Linear mixed-effects model for O-Information across rest segments
The model includes drug condition (placebo, ketamine) as a predictor, with a random intercept for each participant to account for repeated measures. Unstandardized coefficients (Estimates) and standardized beta estimates (std. Beta), along with their 95% confidence intervals (CI) and standard error (SE), are reported.

| | O-Information Moving Experience | | | | | | | |
|---|---|---|---|---|---|---|---|---|
| *Predictors* | *Estimates* | *SE* | *std. Beta* | *std. SE* | *CI (95%)* | *std. CI (95%)* | *t-statistic* | *p-Value* |
| Intercept | 32.53 | 0.67 | 0.02 | 0.11 | 31.20 – 33.85 | -0.19 – 0.23 | 48.28 | **<0.001** |
| Moving | -0.15 | 0.05 | -0.07 | 0.02 | -0.24 – -0.05 | -0.12 – -0.03 | -3.02 | **0.003** |

| | | | | | | | | |
|---|---|---|---|---|---|---|---|---|
| Drug Condition [Placebo] | -0.29 | 0.27 | -0.05 | 0.05 | -0.82 – 0.23 | -0.14 – 0.04 | -1.10 | 0.271 |
| **Random Effects** | | | | | | | | |
| σ2 | 22.30 | | | | | | | |
| τ00 subject | 11.37 | | | | | | | |
| ICC | 0.34 | | | | | | | |
| N subject | 32 | | | | | | | |
| Observations | 1256 | | | | | | | |

Table S6: Linear mixed-effects model for O-Information across moving experiences
The model includes the strength of the moving experience (ranging from 1-10) and drug condition (placebo, ketamine) as predictors, with a random intercept for each participant to account for repeated measures. Unstandardized coefficients (Estimates) and standardized beta estimates (std. Beta), along with their 95% confidence intervals (CI) and standard error (SE), are reported.

| | **3-way Interaction** | | | | | | | |
|---|---|---|---|---|---|---|---|---|
| *Predictors* | *Estimates* | *SE* | *std. Beta* | *std. SE* | *CI (95%)* | *std. CI (95%)* | *t-statistic* | *boot.. p-Value* |
| Intercept | 5.52 | 0.18 | 0.07 | 0.07 | 5.16 – 5.88 | -0.06 – 0.20 | 1.01 | 0.273 |
| Baseline Cluster | -8.75 | 10.13 | 0.02 | 0.07 | -28.63 – 11.13 | -0.13 – 0.16 | 0.24 | 0.790 |
| Hedonic Cluster | 43.31 | 9.36 | 0.13 | 0.05 | 24.94 – 61.68 | 0.04 – 0.22 | 2.82 | **0.012** |
| Drug [Placebo] | -0.46 | 0.22 | -0.11 | 0.07 | -0.90 – -0.02 | -0.24 – 0.02 | -1.62 | **0.036** |
| Baseline Cluster x Hedonic Cluster | -1526.14 | 524.38 | -0.14 | 0.05 | -2554.90 – -497.38 | -0.23 – -0.05 | -2.91 | **0.002** |
| Baseline Cluster x Drug [Placebo] | 11.09 | 11.14 | -0.02 | 0.08 | -10.77 – 32.95 | -0.17 – 0.14 | -0.21 | 0.826 |
| Hedonic Cluster x Drug [Placebo] | -23.48 | 15.03 | -0.03 | 0.07 | -52.97 – 6.01 | -0.16 – 0.10 | -0.47 | 0.455 |
| Baseline Cluster x Hedonic Cluster x Drug [Placebo] | 1820.41 | 580.09 | 0.17 | 0.05 | 682.36 – 2958.46 | 0.06 – 0.27 | 3.14 | **0.002** |

| | |
|---|---|
| **Random Effects** | |
| $\sigma^2$ | 7.40 |
| $\tau_{00\ subject}$ | 0.52 |
| ICC | 0.07 |
| $N_{subject}$ | 32 |
| Observations | 1260 |

Table S7: Linear mixed-effects model predicting hedonic experiences from NBS cluster activity across drug conditions.
The model includes the trial-averaged activity of the hedonic cluster, the run-wise averaged activity of the ketamine baseline cluster, and the drug condition (placebo vs. ketamine) as fixed effects. A random intercept for each participant accounts for repeated measures. Unstandardized coefficients (Estimates) and standardized beta estimates (std. Beta), along with their 95% confidence intervals (CI) and standard error (SE), are reported. Shown p-values were calculated using 10000 wild bootstrap (boot.) iterations.

**Hedonic experience predicted by $\Phi^R$ and rtr**

| *Predictors* | *Estimates* | *SE* | *CI (95%)* | *t-statistic* | *p-Value* |
|---|---|---|---|---|---|
| Intercept | 5.70 | 0.30 | 5.15 – 6.29 | 18.94 | **<0.001** |
| $\Phi^{R+}$ | 19.03 | 7.22 | 4.87 – 33.20 | 2.64 | **0.008** |
| $\Phi^{R-}$ | -36.99 | 12.05 | -60.63 – -13.35 | -3.07 | **0.002** |
| $rtr^+$ | -27.01 | 3.77 | -34.42 – -19.61 | -7.16 | **<0.001** |
| $rtr^-$ | 54.77 | 8.85 | 37.41 – 72.14 | 6.19 | **<0.001** |
| **Random Effects** | | | | | |
| $\sigma^2$ | 6.54 | | | | |
| $\tau_{00\ subject}$ | 0.58 | | | | |
| ICC | 0.08 | | | | |
| N $_{subject}$ | 32 | | | | |
| Observations | 1260 | | | | |

Table S8: Linear mixed-effects model predicting the intensity of the hedonic experience from decomposed NBS hedonic cluster activity.
The model predicts the strength of the hedonic experience using the four decomposed dynamics per trial ($\Phi R^+$, $rtr^+$, $\Phi R^-$, and $rtr^-$). A random intercept for each participant accounts for repeated measures. Parameter estimates are reported along with their 95% confidence intervals (CI) and standard errors (SE). Reported results are based on cluster-robust covariance matrices (CR1).

**Dominance analysis of the effects of $\Phi^R$ and rtr on hedonic experiences**

| *Predictors* | *R2 estimate* | *SE* |
|---|---|---|
| $\Phi^{R+}$ | 0.004 | 0.003 |
| $\Phi^{R-}$ | 0.014 | 0.007 |
| $rtr^+$ | 0.077 | 0.013 |
| $rtr^-$ | 0.052 | 0.012 |

Table S9: Dominance analysis of the linear mixed-effects model predicting the intensity of the hedonic experience from decomposed NBS hedonic cluster activity.
Reported results are based on 1000 bootstrap iterations. Changes in marginal R2 and standard errors (SE) are reported.

**Treatment predicted by $\Phi^R$ and rtr**

| *Predictors* | *Odds Ratios* | *SE* | *CI* | *Statistic* | *p* |
|---|---|---|---|---|---|
| Intercept | 0.94 | 0.10 | 0.76 – 1.16 | -0.57 | 0.570 |
| $\Phi^{R+}$ | 2.10 | 0.27 | 1.64 – 2.73 | 5.69 | **<0.001** |
| $\Phi^{R-}$ | 0.31 | 0.06 | 0.21 – 0.43 | -6.56 | **<0.001** |
| $rtr^-$ | 2.50 | 0.35 | 1.92 – 3.34 | 6.48 | **<0.001** |

| | | | | | |
|---|---|---|---|---|---|
| $rtr^+$ | 0.45 | 0.06 | 0.34 – 0.58 | -6.07 | **<0.001** |
| Observations | 509 | | | | |

Table S10: General linear model predicting treatment from decomposed NBS baseline cluster activity.
The model predicts drug treatment using the four decomposed dynamics per trial ($\Phi R^+$, $rtr^+$, $\Phi R^-$, and $rtr^-$) and a logit-link function. Odds Ratios are reported along with their 95% confidence intervals (CI) and standard errors (SE).

**Dominance analysis of the generalized linear model predicting treatment using $\Phi^R$ and rtr dynamics**

| *Predictors* | *R2 estimate* | *SE* |
|---|---|---|
| $\Phi^{R+}$ | 0.055 | 0.020 |
| $\Phi^{R-}$ | 0.104 | 0.025 |
| $rtr^+$ | 0.072 | 0.024 |
| $rtr^-$ | 0.108 | 0.029 |

Table S11: Dominance analysis of the generalized linear model predicting drug treatment from decomposed NBS baseline cluster activity.
Reported results are based on 1000 bootstrap iterations. Changes in marginal R2 and standard errors (SE) are reported.

**Results of the exploratory receptor associated with the ketamine cluster found during music perception**

Similar to the main analysis, we conducted separate analyses for increasing and decreasing ΦWMS edges. To summarize node-level effects, we collapsed the observed edge dynamics from the NBS analysis by averaging the t-values of each significantly identified edge per node. These node dynamics were then compared with the 19 receptor maps described in the main manuscript.

The results of this exploratory analysis revealed that ketamine-induced increases in $\Phi^{WMS}$ during music perception were significantly and negatively correlated with the density of several neurotransmitter systems, including D2 ($r = -0.43$, $p < 0.05$), DAT ($r = -0.46$, $p < 0.05$), H3 ($r = -0.38$, $p < 0.05$), MOR ($r = -0.40$, $p < 0.05$), and VAChT ($r = -0.42$, $p < 0.05$). Notably, there was also a negative trend between ΦWMS increases and NMDA receptor density ($r = -0.44$, $p = 0.05$). These results are not corrected for multiple comparisons.
